# Rejuvenated human amniotic fluid stem cells: a superior source of standardized induced mesenchymal stem cells for enhanced therapeutic applications

**DOI:** 10.1101/2025.03.25.645040

**Authors:** Michelangelo Corcelli, Ellen Petzendorfer, Kate Hawkins, Filipa Vlahova, Catherine Caruso, Mehedi Mohammad Hasan, Katie Durrant, Anna David, Fleur S van Dijk, Pascale V Guillot

**Affiliations:** University College London, Elizabeth Garrett Anderson Institute for Women’s Health, Research Department of Maternal and Fetal Medicine, London, UK; Department of Development and Regeneration, Katholieke Universiteit Leuven, Belgium; Specialist Neonatal and Paediatric Surgery, Great Ormond Street Hospital NHS Trust, London, UK; Northwest Thames Regional Genetics Service, London Northwest University Healthcare NHS Trust, London, UK; Department of Metabolism, Digestion and Reproduction, Section of Genetics and Genomics, Imperial College London, London, UK

## Abstract

Human fetal mesenchymal stem cells (hfMSCs) present advantageous characteristics compared to their adult counterparts and have emerged as potent cells in the field of regenerative medicine. In the context of skeletal regeneration, human amniotic fluid stem cells (AFSCs) have been shown to improve the quality and structure of the bone extracellular matrix in an experimental model of severe osteogenesis imperfecta. However, primary hfMSCs undergo replicative senescence during in vitro expansion, along with a progressive decrease in plasticity and tissue repair potential. To overcome this challenge, we rejuvenated AFSC to pluripotency using non-integrative episomal reprogramming and subsequently re-derived the cells towards the mesoderm to obtain induced MSCs (iMSCs). We found that iMSCs have a slower proliferation rate compared to their parental cell line (40h±2h vs. 29h±5h) but retain the multipotency and differentiation potential characteristic of MSCs. Comparative genomic analysis revealed that iMSCs express higher levels of genes involved in maintaining stemness, cell signaling, adhesion and migration, as well as promoting osteoblast differentiation, whilst AFSC expressed higher levels of genes involved in cell proliferation. In addition, iMSCs secrete small extracellular vesicles (iEVs) that have the potential to stimulate fibroblast migration, a key process in tissue repair and wound healing. Together, these data suggest that resetting the epigenetic clock of primary hfMSCs may represent a promising strategy to address the limitations associated with primary cell use and enhance their therapeutic potential.

## Introduction

Human mesenchymal stem cells (MSCs) have emerged as a promising tool in regenerative medicine due to their multipotent differentiation capacity, immunomodulatory properties, and ability to promote tissue repair. These cells can differentiate into mesodermal lineages, including osteocytes, chondrocytes, and adipocytes, making them valuable candidates for bone regeneration, cartilage repair, and other clinical applications. While adult MSCs derived from bone marrow, adipose tissue, and other sources have been extensively studied, fetal mesenchymal stem/stromal cells (hfMSCs) isolated during pregnancy from various fetal tissues^1^, placenta^2^ and amniotic fluid^3,4^ have attracted growing interest due to their enhanced proliferative capacity, superior differentiation potential, and reduced immunogenicity compared to their adult counterparts.^5–7^ Human fetal MSCs, particularly those derived from amniotic fluid, possess several advantages over adult MSCs. They exhibit higher telomerase activity, longer telomeres, and increased plasticity, allowing for more robust expansion in vitro.^7^ These characteristics make fetal MSCs attractive for therapeutic applications. Indeed, the regenerative potential of primary hfMSCs isolated during the first trimester of pregnancy has been reported in various preclinical models, including Alport disease^8^, brittle bone disease^9–13^ and neonatal encephalopathy^14^. The regenerative potential of fetal tissue-derived hfMSCs has also been investigated in clinical trials.^15,16^

Human mid-trimester amniotic fluid, which is accessible during ongoing pregnancy, has been identified as an alternative source of potent fetal stem cells (amniotic fluid stem cells or amniocytes, AFSCs)^3,17^ for cell therapy, tissue engineering and disease modelling applications.^3,4^ AFSCs are easy to reprogram to pluripotency and were shown to be receptive to chemically-induced reprogramming using small molecule epigenetic modifiers.^18–20^ Human AFSCs have also emerged as an effective cell source for skeletal applications.^21,22^ When transplanted perinatally into a mouse model of severe osteogenesis imperfecta, AFSCs showed the potential to counteract bone fragility, reducing fracture susceptibility, increasing bone strength, improving bone quality and micro-architecture, normalising bone remodelling and promoting endogenous osteogenesis and maturation of resident osteoblasts.^13^

However, some hurdles remain before primary MSC can be used routinely in the clinic. First, the properties and therapeutic efficacy of primary hfMSCs depend on their tissue of origin, stage of development, number of *in vitro* cell division and culture conditions.^1,6,23^ Additional caveats include the lack of protocols for the generation of standardized cell preparations, safety concerns, ectopic engraftment, differentiation, immunological rejection and ethical concerns. Finally, the need to obtain a sufficient number of donor cells requires extensive *in vitro* expansion. As MSCs divide repeatedly *in vitro*, they eventually enter a state of replicative senescence, where they lose the ability to proliferate.^24^ This limits the number of cells that can be expanded, making it challenging to obtain sufficient quantities for therapeutic use and clinical application, especially for high-dose or repetitive-dose treatments.^24^ In addition, senescent MSCs show a reduced ability to differentiate into key lineages (osteogenic, chondrogenic, and adipogenic) and may also impact their paracrine efficacy, i.e. their ability to modulate and normalise target cell behaviour. In addition, the ability of MSCs to home to injury sites and engraft into tissues declines with senescence, impairing their capacity to support tissue repair. This represents an issue of standardization, as new donors need to be identified repeatedly to isolate new hfMSCs batches, which constitutes a barrier to clinical implementation and introduces variability of therapeutic intervention outcomes.

To overcome these challenges, human induced pluripotent stem cells (iPSCs) have emerged as a powerful alternative source of MSCs. Generated by reprogramming somatic cells through the forced expression of key transcription factors (such as OCT4, SOX2, KLF4, and c-MYC), iPSCs can proliferate indefinitely while maintaining the potential to differentiate into cells of all three germ layers. Importantly, iPSCs can be further directed to generate MSC-like cells, known as induced MSCs (iMSCs), which exhibit similar phenotypic and functional characteristics to primary MSCs. This approach offers an unlimited source of patient-specific or allogeneic MSCs, circumventing the issue of donor-dependent variability and overcoming the limitations of replicative senescence.^25–28^ This cost-effective strategy enables to achieve consistent, reproducible iMSCs manufacture at commercial scale whilst maintaining optimal therapeutic potential and eliminating the need for multiple donors.

In this study, we reprogrammed human mid-trimester AFSCs into iPSCs and subsequently differentiated these iPSCs into iMSCs. This strategy not only preserves the advantages characteristics of fetal MSCs but also provides a renewable source of MSCs capable of sustained expansion and clinical scalability. Our objective was to establish a robust protocol for deriving iMSCs from human AFSC, characterize their molecular and functional properties, and compare them to their parental MSC counterparts. By leveraging the proliferative advantages of iPSCs and the regenerative potential of hfMSCs, this work aims to develop a clinically relevant source of MSCs for future therapeutic applications. Ultimately, this approach may address current limitations in MSC-based therapies and pave the way for more effective and scalable cell-based interventions.

## Material and Methods

### Cell culture

Human mid-trimester amniotic fluid mesenchymal stem cells (AFSCs) (passage 7-10), were isolated from the amniotic fluid of a healthy pregnancy at 12 weeks of gestation, which was obtained under ultrasound guidance and according to the ethical approval given by the Research Ethics Committees of Hammersmith & Queen Charlotte’s Hospitals (08/H0714/87) in compliance with national guidelines (Polkinghorne) for the collection of fetal tissue for research, as we previously described.^13^ In brief, the cells from three individuals were selected for c-KIT expression and expanded at a density of 10^4^cells/cm^2^ on plastic culture dishes without feeders in Dulbecco’s modified Eagle’s medium (DMEM-HG) (Thermo Fisher) supplemented with 10% fetal bovine serum (Biosera), 2 mM L-glutamine, 50 IU/ml penicillin and 50mg/ml streptomycin (Thermo Fisher), at a density of 1×10^4^ cells/cm^2^ at 37°C in a 5% CO_2_ incubator.

Human primary dermal fibroblast isolated from neonatal foreskin (PCS-201-010, LGC Standards) were cultivated in fibroblast basal medium containing essential and non-essential amino acids, vitamins, other organic compounds, trace minerals, and inorganic salts (CS-201-03, LGC Standards).

### Derivation of iMSCs from human AFSC-derived iPSCs

AFSC were reprogrammed to pluripotency using non-integrative episomal reprogramming, following the manufacturer’s instructions (A15960, Thermo Fisher). In short 500,000 AFSC were dissociated with TrypLE and resuspended in 90ul electroporation buffer supplemented with 20 ul of hSK, hUL and hoct4. The cells were electroporated using an electroporator (Neon Transfection System, Thermo Fisher), programme C17 and plated in AFSC expansion medium onto a Geltrex-coated vessel (Thermo Fisher) vessel at 37°C in a 5% CO_2_ incubator. Upon reaching confluence the next day, the cells were detached and reseeded onto a Geltrex-coated in in Essential E8 culture medium (both from Stemcell Technologies). Colonies were picked on day 20 using a p200 pipette tip, pooled and reseeded in E8 medium supplemented with ROCK inhibitor (Thermo Fisher).

When reaching 50% confluency, AFSC-derived iPSCs were detached using ReLeSR (Stemcell Technologies) and replated as single cells in Essential E8 culture medium. The following day, the culture medium was replaced by mesoderm induction medium, and replaced every day for the following 4 days, upon which the medium was replaced by L-glutamine-supplemented mesencult medium (Stemcell Technologies). The mesencult medium was replaced every 48 hours. On day 10, the cells were detached using TrypLE Express (Thermo Fisher) and reseeded at a density of 5×10^4^ cells/cm^2^ on attachment substrate-coated plastic vessels in mesencult medium. Once the cells had reached 70% confluency, they were detached and reseeded at a density of 3×10^4^ cells/cm^2^. At passage 3, the cells were reseeded on attachment substrate-coated plastic vessels in mesencult medium at a density of 1×10^4^ cells/cm^2^. The cells were cultivated at 37°C in a 5% CO_2_ incubator.

### *In vitro* differentiation

Cells were differentiated along the osteoblast lineage for 3 weeks in DMEM-LG supplemented with 10 mM β-glycerophosphate, 0.2 mM ascorbic acid and 10^-8^M dexamethasone, then fixed in 10% formalin. Cells were differentiated along the adipocyte lineage over 2 weeks in DMEM supplemented with 0.5 mM hydrocortisone, 0.5 mM isobutyl methylxanthine and 60 mM indomethacin, then fixed and stained with oil red O. Cells were differentiated along the chondrocyte lineage over 3 weeks in DMEM-LG supplemented with 0.01 µg/ml TGF-β3, 0.1 µM dexamethasone, 0.17mM ascorbic acid, 1 mM sodium pyruvate, 0.35 mM L-proline, 1% ITSS, 50 µg/ml linoleic acid (all reagents from Merck), then cells were fixed in and stained with Alcian blue (2%).

### Flow cytometry

The cells were detached, washed in flow buffer (PBS+3%BSA, Merck) and centrifuged at 5000g for 2 minutes before 1×10^5^ cells were resuspended in the appropriate primary antibody, i.e. CD73-PE, CD90-APC and CD105-FITC (Biolegend) at 1:10 dilution in flow buffer and incubated for 1 hour at 4°C. The cells were then washed again with the FACS buffer and analysed using a Becton Dickinson FACScalibur flow cytometer (BD Biosciences) using Cell Quest Pro and FlowJo software. Negative controls were unstained cells.

### Immunofluorescence

The cells were washed in PBS, fixed in 4% paraformaldehyde (PFA, Merck) and permeabilized using Triton X-100 (Merck). The cells were then blocked for 30 minutes with blocking buffer (PBS supplemented with 2% bovine serum albumin and 0.1% Tween) and incubated overnight with primary antibodies at their optimal dilution, i.e. OCT4A (Santa Cruz), KLF4 (Stemgent), SOX2 (Abcam), NANOG (Miltenyi Biotec), TRA-1-60, (Abcam), CD90 (Abcam), CD73 (BD Biosciences), CD105 (BD Biosciences), CD34 (Dako Cytomation), CD45 (Dako Cytomation), then washed and incubated with secondary antibody (Alexa Flour 488 Goat anti-rabbit IgG, Alexa Flour 488 Goat anti-mouse IgG, all from Abcam 1:500) for 1 hour at room temperature. Cells were then counter-stained with 4’,6-diamidino-2-phenylindole (DAPI) and visualized immediately. Live cell imaging was performed using TRA-1-60 Alexa Fluor 488 conjugated kit, following the manufacturer’s instructions (Thermo Fisher Scientific). Images were collected using a Leica DM 6000 fluorescence microscope (40x PLAN APO objective) and transferred to Adobe Photoshop (Adobe Systems).

### Growth kinetics

Growth kinetics were assessed for each cell type (iMSCs and AFSCs) in triplicates at 10^4^ cells/cm^2^ in 10cm plates in growth medium. Cells were detached when sub-confluent and counted in trypan blue to exclude dead cells in a hemocytometer before replating in similar density. The doubling times were calculated via this formula: DT=t/(Log_2_[y/m]), with DT=doubling time, t=time in culture, y=number of cells at end of culture, m=number of cells at beginning of culture, as described previously^7^.

### Gene expression analysis

Total RNA was extracted from 8-week-old mice femoral epiphysis, using TRIzol (Thermo Fisher Scientific), followed by RNA clean-up (RNeasy Qiagen) and cDNA synthesis using RT^2^ First Strand Kit (Qiagen, Germany). Gene expression was examined by RT^2^ Profiler human PCR arrays (PAHS-082Z, Qiagen) and analysed according to the manufacturer’s instructions (n=3 per group).

### Isolation and characterisation of extracellular vesicles (iEVs)

Size exclusion chromatography (SEC) was used to isolate iEVs from the conditioned culture supernatant of iMSCs. The iMSCs culture supernatant was collected and subjected to differential centrifugation, i.e. 300×g for 10 minutes, followed by 2000×g for 10 minutes, and 20,000×g for 30 minutes. The supernatant was then concentrated to 500-1000 µL using 10 kDa Amicon Ultra centrifugal filter units (Merck) before iEVs isolation. A commercially available size exclusion chromatography (SEC) column qEVoriginal/70 nm (Izon Sciences) was used for the isolation of EVs by following the standard manufacturer protocol. In brief, the column was prewashed with 17 mL filtered (0.2 µm Minisart syringe filters) Dulbecco’s phosphate-buffered saline (DPBS, Merck Life Science) and 500 µl of the concentrated sample was added to the top of the column filter. After the sample had passed down, 6 mL of DPBS was added immediately on the top of the column filter. The first 2.5 ml were discarded, and the 5 following fractions (each 500 µl) were pooled and concentrated using 10 kDa Amicon Ultra 2 ml centrifugal filter units. The size profile and concentration of iEVs were then determined using Tuneable Resistive Pulse Sensing (TRPS), using a 150 nm nanopore calibrated with CPC200 particles (NIST-traceable carboxylated polystyrene, 200 nm), and PBS as electrolyte (Izon Sciences).

### Transmission Electron Microscopy (TEM) imaging of iEVs

iEVs suspension was deposited on Formvar-carbon-coated 200 mesh copper grids (Agar Scientific, Stansted, UK) and fixed using 2% paraformaldehyde (Merck Life Science, UK) with 1% glutaraldehyde (Polysciences). The vesicles were contrasted in uranyl oxalate and embedded in a mixture of methylcellulose (Merck) and uranyl acetate (Polysciences). Samples of iEVs were then observed using Phillips/FEI CM 120 BioTwin TEM at 120 kV for imaging.

### Cellular uptake of iEVs

iEVs were incubated with BioTracker 655 Red Cytoplasmic Membrane Dye (SCT108-Merck) for 15 minutes at RT at a dilution of 1:200 before being subjected to two steps of centrifugal filtration of 20 minutes at 4000g at +4°C using a 100 kDa MWCO 0.5ml Amicon ultra centrifugal filter (Merck) to remove any unbound dye. Labelled iEVs were then resuspended in DMEM basal culture medium and administered to fibroblast cells. Cells were visualised after 2 hours with fluorescence microscopy using a Zeiss Axio Observer A1 inverted microscope. Negative controls with no dye and positive controls with dye only were also acquired.

### *In vitro* migration assay

5×10^4^ human fibroblasts were seeded into each side a removable 2 well culture insert in a µ-Dish 35 mm plate, to create a cell-free gap for migration assays (Ibidi). After removing the insert, iEVs were added to the growth medium (DMEM supplemented with 5% EV-depleted FBS) at a concentration of 10^8^ EVs/ml and the gap was monitored at 0 hour, 24 hours, 48 hours and until complete closure. Phase contrast images were acquired using a Zeiss or Leica inverted microscope using a 5x objective. Gap width and cell-free area were measured using the open-source ImageJ wound healing tool (Fiji). The assay was used to assess the effects of novel therapeutics like EVs in increasing the migration/proliferation of fibroblasts *in vitro*.

## Results

### Mesodermal differentiation of human iPSC into iMSC

To derive iMSCs, human iPSCs from a healthy donor were initially plated on Geltrex matrix in essential 8 medium, which is a chemically-defined serum-free, xeno-free cell culture medium that supports long-term pluripotency. The cells formed distinct clusters of colonies (**Figure 1A**) which expressed markers associated with pluripotency such as TRA-1-60 (**Figure 1B**), Nanog, OCT4A, SOX2, and KLF4 (**Figure 1C**). The cells were subsequently detached and replated as single cells on Geltrex matrix in essential 8 medium, with the medium being replaced by mesoderm induction medium the following day, to induce early mesoderm differentiation over a 5-day period (**Figure 1D**). The cells were further differentiated down the mesenchymal lineage by replacing the medium with L-glutamine-supplemented mesencult medium, which is animal component-free (ACF), serum-free and extracellular vesicle (EV)-free, and plated on attachment substrate-coated plates. The cells were passaged at 70% confluency and a 1 to 3 ratio for 3 weeks, upon which they were characterised.

**Figure 1.**
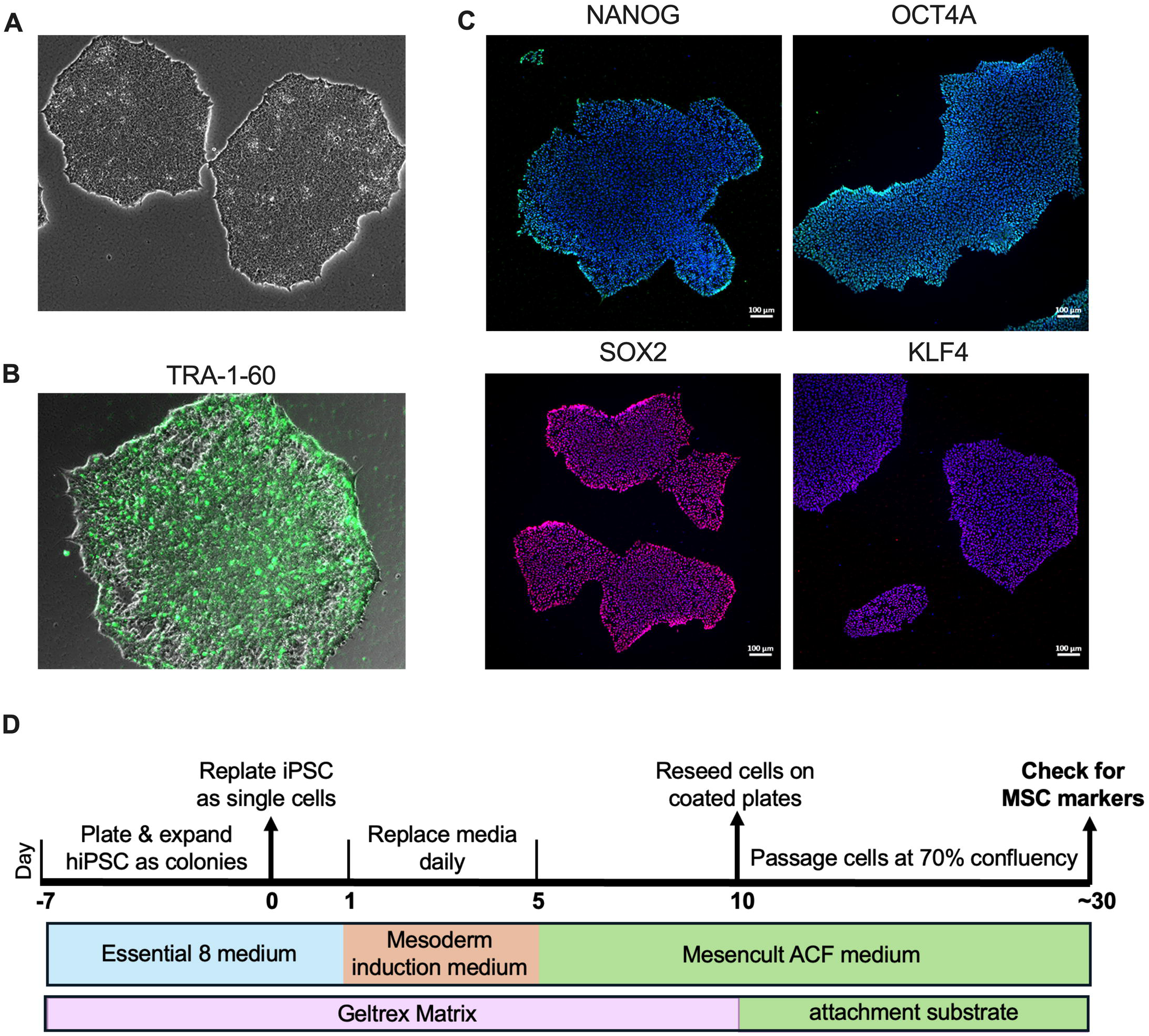
Differentiation of hiPSCs into iMSCs. **A.** Schematic representation of the differentiation of hiPSCs into iMSCs. **B.** Phase contrast image of hiPSCs (original magnification is x40). **C.** Immunofluorescence showing expression of TRA-1-60 in live hiPSCs (green), (original magnification is x40). **D.** Immunofluorescence showing expression of the pluripotency-associated markers NANOG (green), OCT4A (green), SOX2 (red), and KLF4 (red), nuclei were counterstained with DAPI (blue). Scale bar is 100 µm.

### Human iPSC-derived iMSCs present a similar phenotype to AFSCs

Phenotypically iMSCs have a spindle shape morphology and grow as single cells on plastic (**Figure 2A**). Growth kinetic analysis over 3 weeks revealed that iMSC have a doubling time of 40h±2h, which is significantly slower than the 29h±5h of their parental AFSC line (mean±SD, P<0.05) (**Figure 2B**). The cells are negative for OCT4A and express vimentin, with 100% cells co-expressing the mesenchymal-associated markers CD73, CD90 and CD105, as assessed by immunofluorescence and flow cytometry (**Figure 2C, D and E**). The cells are negative for the hematopoietic markers CD34 and CD45 (**Figure 2D**). When cultivated in chondrogenic, adipogenic and osteogenic permissive conditions, the cells can differentiate down the three lineages, as assessed by Alcian blue staining of collagen (chondrogenic), oil red o staining of lipid droplets (adipogenic) and alizarin red staining of mineral deposition (osteogenic) (**Figure 3A and 3B**).

**Figure 2.**
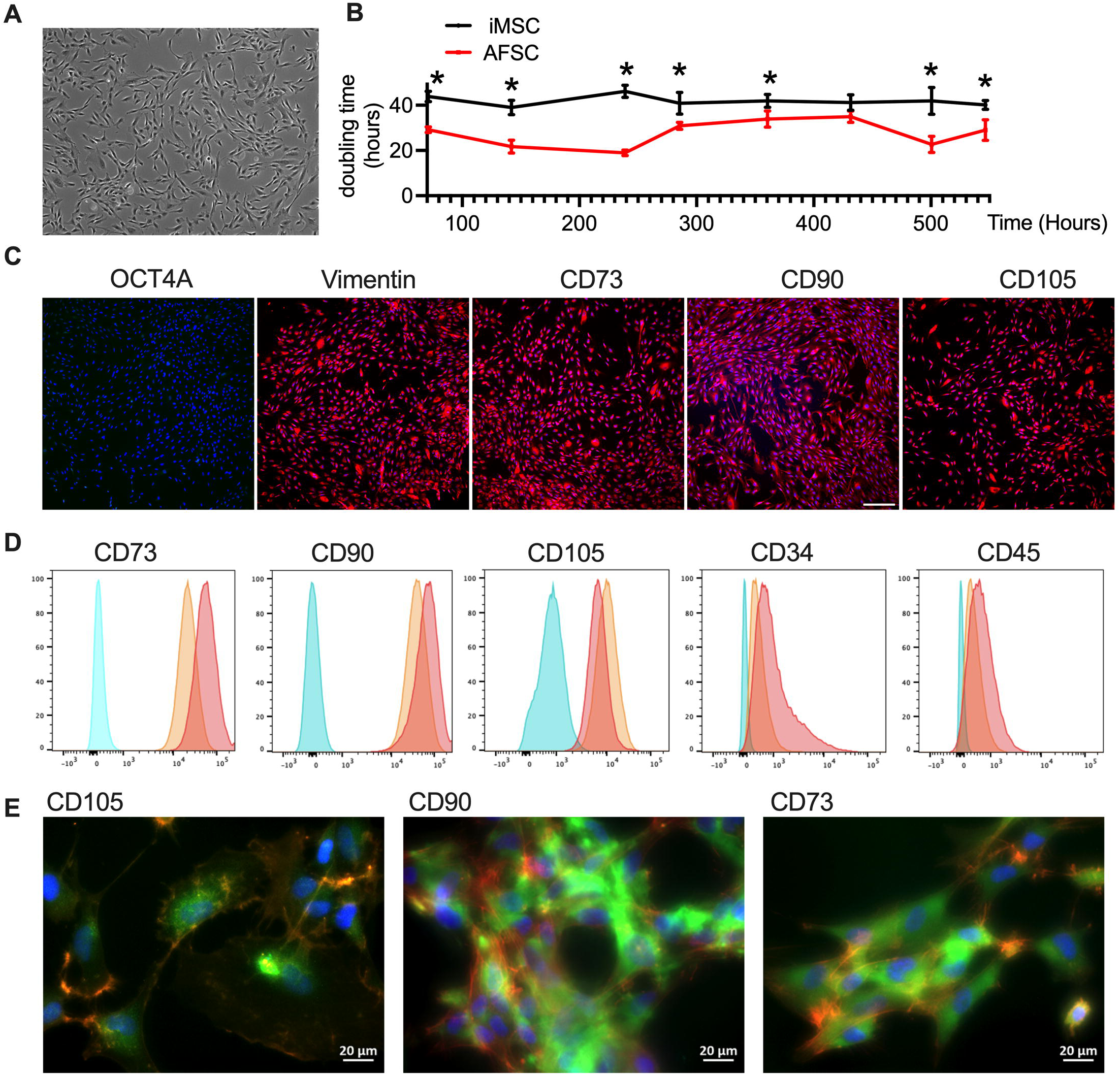
Characterisation of iMSCs. **A.** Phase contrast image of iMSCs (original magnification is x40). **B.** growth kinetics of iMSCs and AFSCs over 600 hours in culture in expansion media. **C.** Immunofluorescence showing lack of expression of the pluripotency-associated marker OCT4A (green) and expression of the MSC-associated markers vimentin, CD73 (red), CD90 (red) and CD105 (red), nuclei were counterstained with DAPI (blue) (original magnification 40X). **D.** CD73, CD90, CD105, CD34 and CD45 were assessed by flow cytometry, the red tracing shows stained iMSCs, the orange tracing shows stained AFSC (positive control), and the blue tracing shows unstained iMSCs (negative control). **E.** Immunofluorescence showing expression of the MSC-associated markers CD105 (green), CD90 (green) and CD73 (green), nuclei were counterstained with DAPI (blue), actin filaments were counterstained with phalloidin (red). Scale bar is 20 µm.

**Figure 3.**
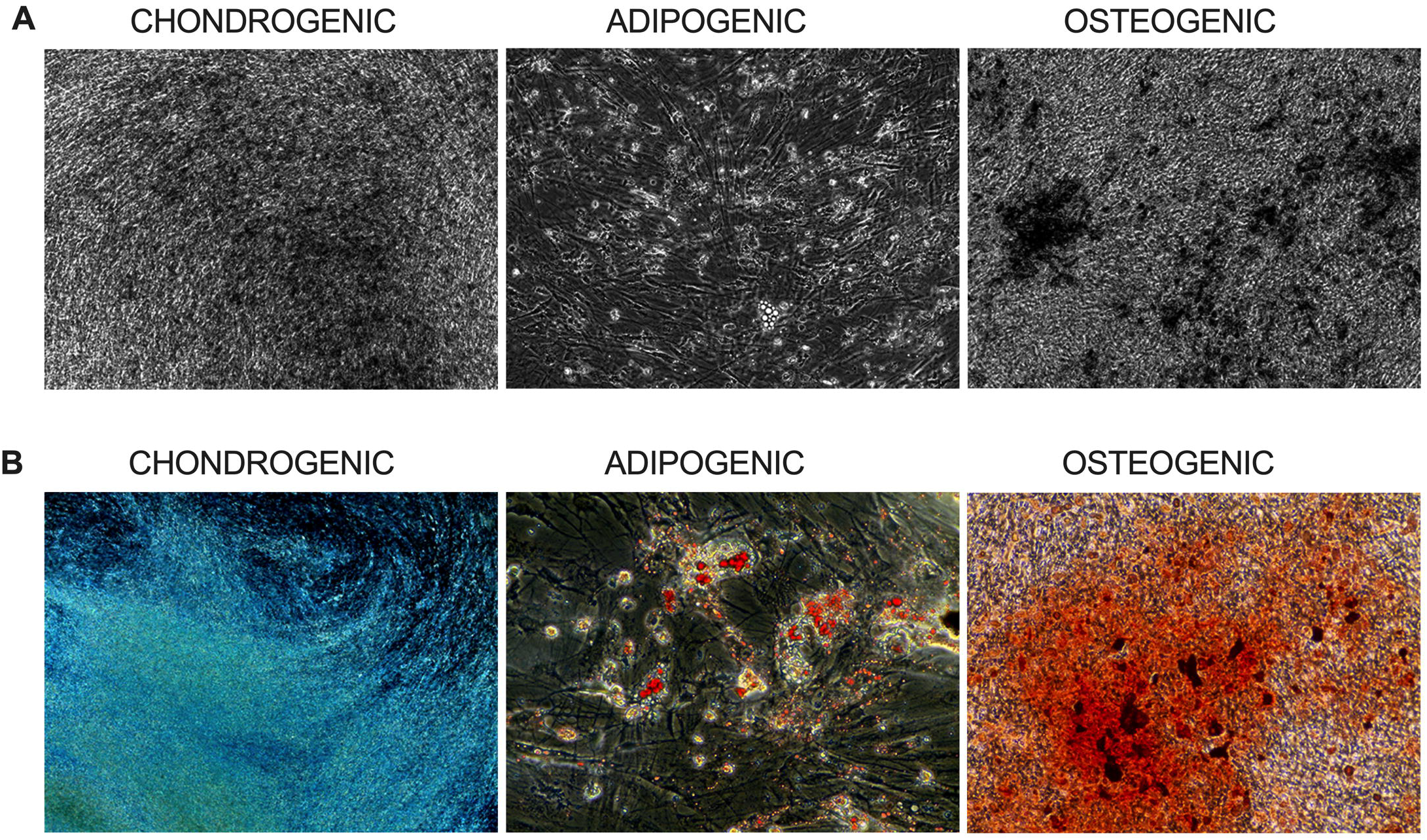
Tri-lineage differentiation potential of iMSCs. **A.** Phase contrast images of iMSCs after 21 days in chondrogenic medium (chondrogenic), 14 days in adipogenic medium (adipogenic) and 21 days in osteogenic medium (osteogenic), original magnification x20. **B.** Collagen visualised using Alcian blue staining (chondrogenic), lipid droplets stained with Oil red O (adipogenic) and minerals stained with Alizarin red staining (osteogenic), original magnification x20.

### iMSCs and AFSCs express a different set of genes

An RT^2^ PCR profiler array of human mesenchymal stem cells genes (Qiagen) was used to compare the genetic profile of iMSCs and AFSCs (**Figure 4**). Out of the 84 target genes tested in triplicate, 32 genes were differentially expressed, showing at least a two-fold difference in gene expression between both lines. Compared to AFSCs, iMSCs showed upregulation in 27 genes (**Figure 4A**), i.e. ANPEP (Alanyl (membrane) aminopeptidase), CASP3 (Caspase 3, apoptosis-related cysteine peptidase), CTNNB1 (Catenin (cadherin-associated protein), beta 1, 88kDa), ENG (Endoglin), GDF15 (Growth differentiation factor 15), JAG1 (Jagged 1), KAT2B (K(lysine) acetyltransferase 2B), MCAM (Melanoma cell adhesion molecule), PDGFRB (Platelet-derived growth factor receptor, beta polypeptide), PIGS (Phosphatidylinositol glycan anchor biosynthesis, class S), SLC17A5 (Solute carrier family 17 (anion/sugar transporter), member 5), SMAD4 (SMAD family member 4), TGFB1 (Transforming growth factor, beta 1), TGFB3 (Transforming growth factor, beta 3), VCAM1 Vascular cell adhesion molecule 1, ALCAM (Activated leukocyte cell adhesion molecule), ANXA5 (Annexin A5), CD44 (CD44 molecule), COL1A1 (Collagen, type I, alpha 1), FGF2 (Fibroblast growth factor 2 (basic)), HAT1 (Histone acetyltransferase 1), ITGAV (Integrin, alpha V (antigen CD51), ITGB1 (Integrin, beta 1), MMP2 (Matrix metallopeptidase 2), NT5E (5’-nucleotidase, ecto, CD73), THY1 (Thy-1 cell surface antigen) and VEGFA (Vascular endothelial growth factor A). A total of 5 genes were downregulated in iMSCs compared to AFSCs (**Figure 4B**), i.e. CSF3 (Colony stimulating factor 3), EGF (epidermal growth factor), INS (insulin), KITLG (KIT ligand), and HAT1 (Histone acetyltransferase 1).

**Figure 4.**
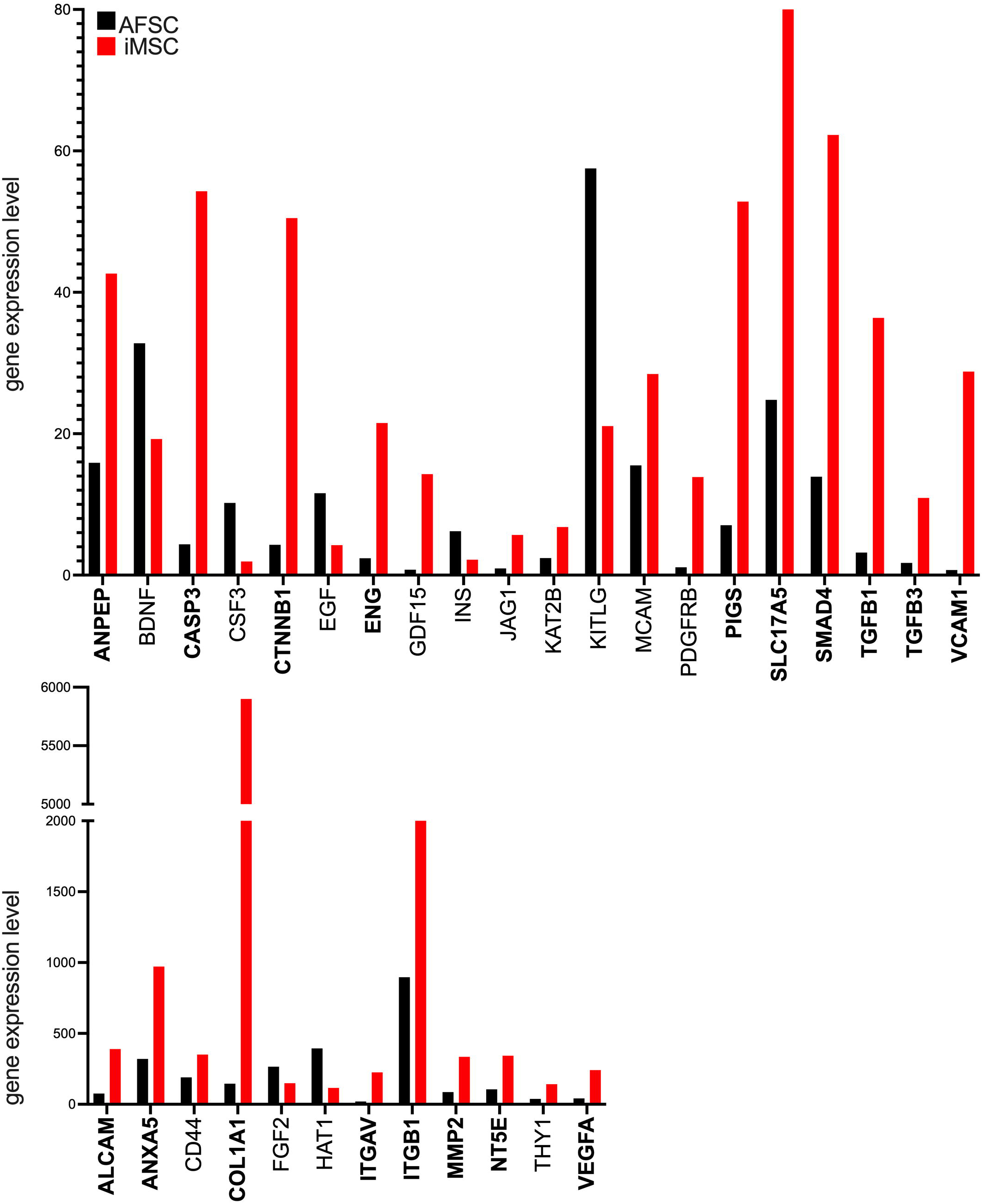
Comparative genomics between iMSCs and human amniotic fluid stem cells (AFSC). Gene expression in iMSCs (red bars) and AFSCs (black bars) using RT2 profiler PCR array PAHS-082Z for human mesenchymal stem cells genes, expressed as 2e-ΔCT, relative to the housekeeping genes GAPDH, ACT2, B2M and HPRT1.

### iMSCs secrete small extracellular vesicles (iEVs) capable of stimulating fibroblast migration *in vitro*

TEM analysis revealed that the extracellular vesicles released by iMSCs in the culture medium (iEVs) exhibit a predominantly spherical (cup-shaped) morphology, consistent with the typical appearance of extracellular vesicles and microvesicles under electron microscopy. The vesicles displayed a well-defined lipid bilayer, further confirming their membranous structure. Minimal aggregation was observed (**Figure 5A**). Delfia analysis showed the expression the tetraspanins CD9, CD81 and CD63 (Figure 5B). The iEVs size range and concentration were determined using Tuneable Resistive Pulse Sensing (TRPS), which measures the change in electrical resistance as individual particles pass through a nanopore. This enables to provide direct particle size and concentration measurement, unlike nanoparticle tracking analysis (NTA, which estimates size based on particle movement/Brownian motion). Analysis showed an average iEV particle diameter range of 96±27.3 nm (mode 78 nm, maximum 238 nm, minimum 68 nm) and a total particle concentration of 2.76e+10 particles/ml.

**Figure 5.**
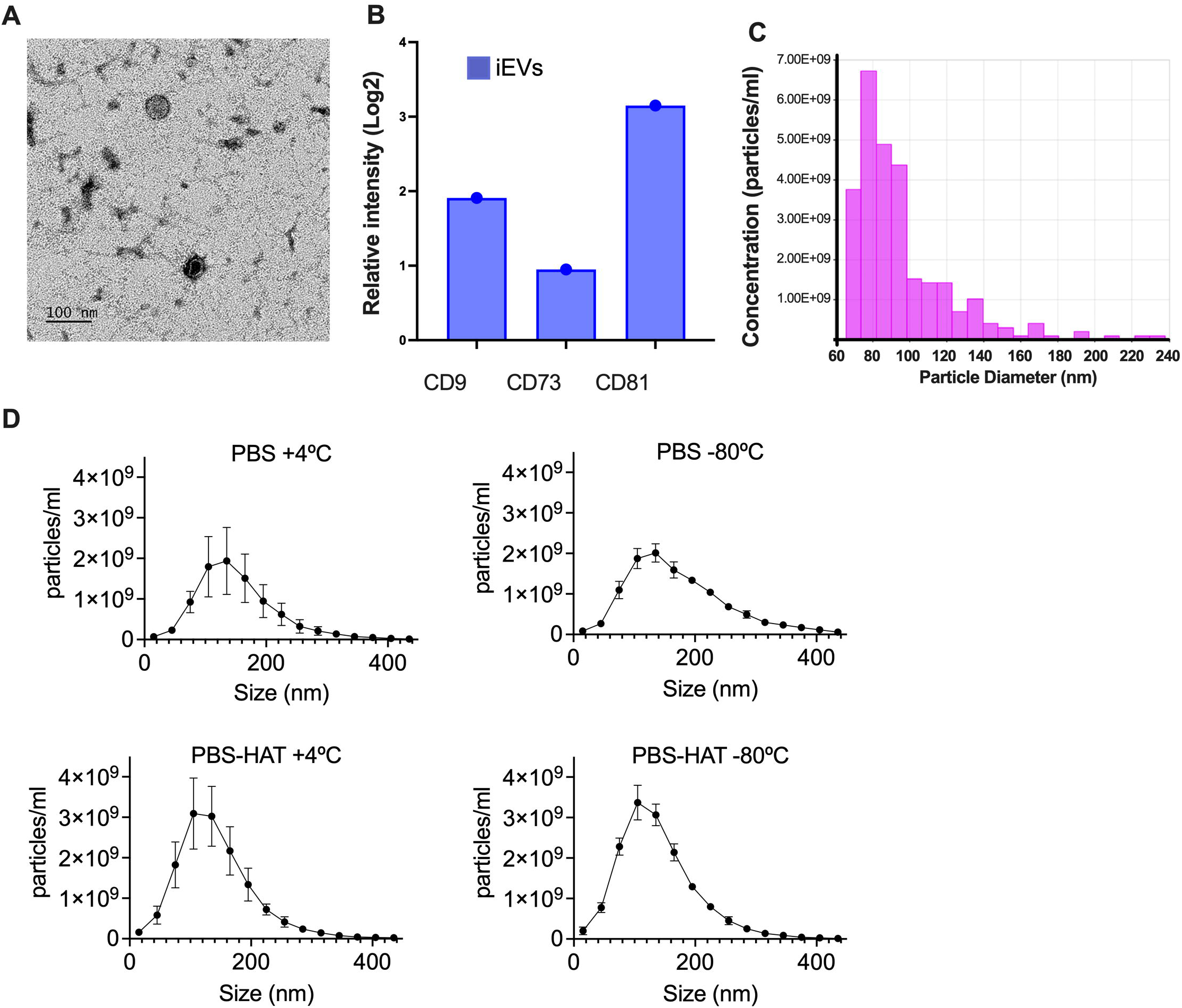
Characterisation of extracellular vesicles secreted by iMSCs (iEVs). **A.** Transmission electron microscopy (TEM) images of iEVs, scale bar is 100 nm. **B.** Levels of tetraspanin EV-specific makers CD9, CD73 and CD81 using DELFIA. **C.** Size distribution of iEV using NanoFCM. **D.** Particle concentration and size distribution of iEVs stored in different conditions for 26 days, i.e. PBS +4°C, PBS - 80°C, PBS-HAT +4°C, PBS-HAT −80°C (mean±SD, n=3).

Comparative analysis of the impact of iEVs storage conditions was assessed after 26 days of storage, based on particle concentration. Results showed that storage at - 80°C in siliconized vessels in PBS supplemented with HEPES 25 mM, human Albumin 0.2% and Trehalose 25 mM (PBS-HAT) led to higher iEVs concentration. Samples stored at +4°C showed larger standard deviations (SDs), indicating greater variability across the particle size distribution (PSD), possibly due to particle aggregation or degradation. In contrast, smaller SDs were observed when stored at - 80°C.

### iEVs stimulate human fibroblast migration *in vitro*

The *in vitro* exclusion assay was used to investigate the potential of iEVs to accelerate human fibroblast migration, as evidence of iEV functionality. The addition of iEVs to human fibroblasts cell culture medium facilitated the migration of cells, as evidenced in the wound healing assay over a 96-hour period (**Figure 6A**). Specifically, after 24h of iEVs treatment fibroblasts had covered 17.86% of the total gap surface) whilst non-treated fibroblasts have not started closing the gap. At 48h, 72h and 96h, non-treated fibroblasts closed 21.43%, 42.86% and 92.14%, respectively. In contrast, iEVs treatment led to a 79.29%, 97.54% and 99.21% gap closure, respectively, for these time points (**Figure 6B**). Fibroblast cell uptake assay revealed that iEVs are taken up into target fibroblast cell body to deliver their cargo intracellularly (**Figure 6C**).

**Figure 6.**
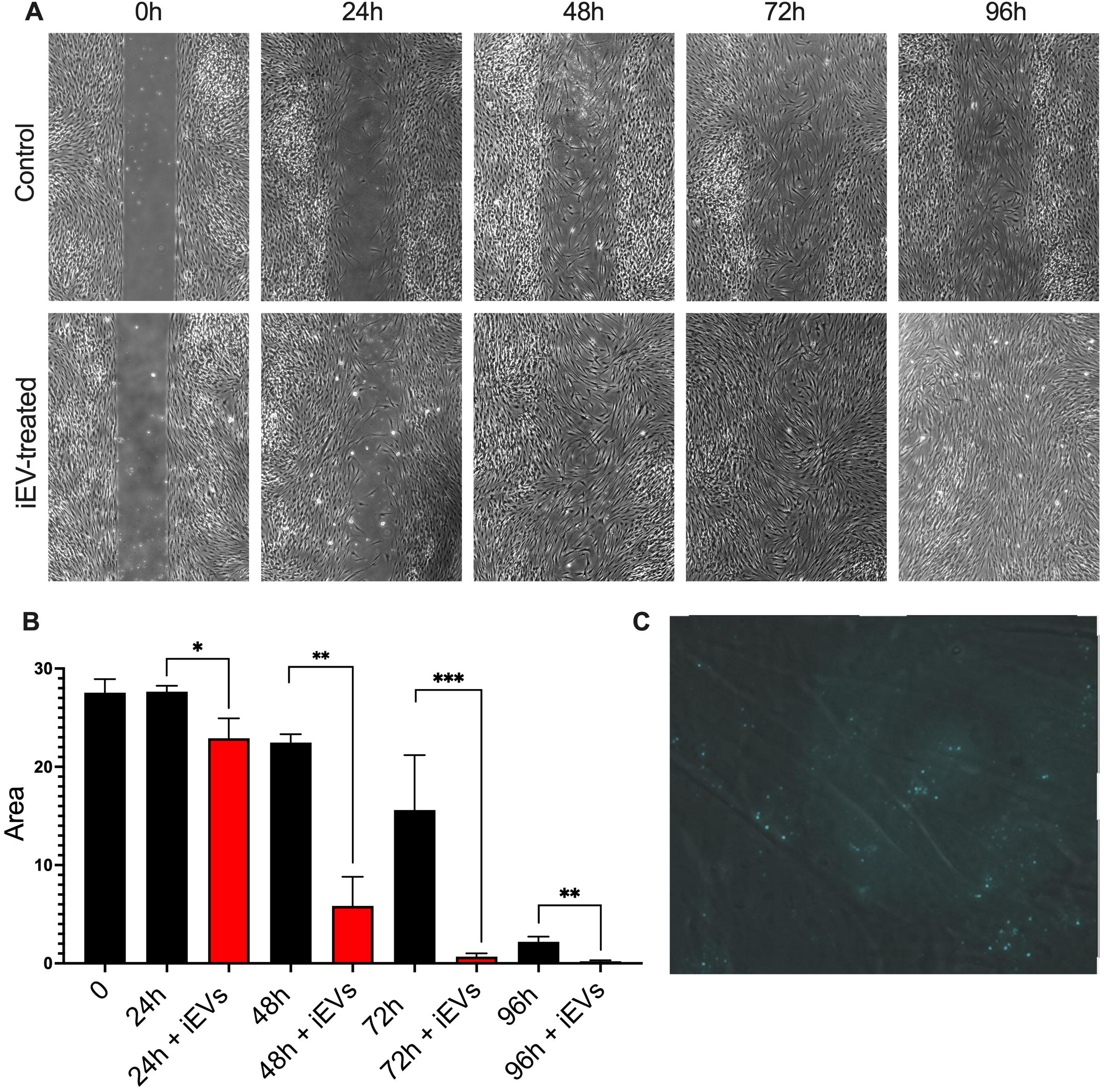
Potential of iEVs to stimulate human fibroblast migration in vitro. **A.** Phase contrast images of human fibroblast cultures just after gap creation using Ibidi insert at 0h, and 24h, 48h, 72h and 96h later in the absence (control) or presence of iEVs (iEV-treated). Original magnification is x20. **B.** Quantification of wound using Fiji, n=3, mean + SD, * P<0.05, ** P<0.01, ***P<0.001. **C.** Visualisation of iEVs labelled with Biotracker Red inside human fibroblasts.

## Discussion

In this study, we successfully derived mesenchymal stem cells (iMSCs) from human iPSCs reprogrammed from AFSCs. Characterisation of iMSCs revealed that the cells present a similar phenotype compared to their primary parental AFSC line. This includes a spindle-shaped morphology, with over 90% cells co-expressing the MSC-associated markers CD73, CD90 and CD105, with absence (or low) expression of CD45 and CD34. The cells retained the multipotency and differentiation potential characteristic of MSCs, i.e. over 90% of cells co-expressing CD73, CD105 and CD90 and being able to differentiate down the osteogenic, adipogenic and chondrogenic pathways. They show a stable proliferation rate over 4 weeks in culture, with a doubling time averaging approximately 41 hours, which is slower than the 28 hours reported for their parental AFSC line. However, this is shorter to that reported for some lines of adult MSCs, beyond passage 6-7.^29^, albeit MSC proliferation rate being dependent if various factors including culture media composition, seeding density and donor age and tissue origin.

At the genetic level, compared to the parental AFSC line, iMSCs exhibited increased expression of genes involved in promoting osteogenic differentiation, as well as genes implicated in the regulation of mesenchymal stem cells (MSCs) proliferation and migration. These molecular changes contribute to modulating their regenerative potential. For example, iMSCs expressed higher levels of expression for ANPEP, CTNNB1, CASP3 and ENG (CD105) genes, which are involved in promoting osteoblast differentiation and regulating osteoclastogenesis. TGFB1 stimulates MSCs differentiation into osteoblast progenitors whilst TGFB3 regulates MSCs migration to injury sites. They also expressed higher levels of SMAD4, which is involved in triggering osteoblast differentiation, as well as ITGB1, PDGFRB, ANXA5, CD44, COL1, which play an important role in regulating proliferation, migration, and differentiation of MSCs, which is important for their ability to home to sites of injury or inflammation. We found that iMSCs also expressed higher levels of genes involved in cell signaling, adhesion, migration and signaling, such as ITGAV, MMP2, PIGS. In addition, iMSCs expressed higher levels of ALCAM, which is also involved in maintaining stemness and keeping MSC in a quiescent state until their receive differentiation signals, as well as CD73, which converts AMP (adenosine monophosphate) into adenosine, and VEGFA, which supports the vascular niche, helping MSCs access oxygen and nutrients, as well as recruiting progenitor cells to sites of injury, contributing to tissue regeneration. In contrast, AFSC expressed higher levels of genes involved in the response to inflammatory signals (CSF3), in cell survival and proliferation (INS and KITLG) and maintenance of the cells in their undifferentiated state (HAT1). Together, these data suggest that iMSCs may be better suited than AFSC for therapeutic applications, in particular skeletal regeneration.

MSCs mediate some of their therapeutic paracrine effects through the extracellular vesicles (EVs) they release. We isolated and characterised the EVs secreted by iMSCs (iEVs). Results showed that iEVs, which secreted in large quantities and have an average diameter size of 96 nm, stimulated fibroblast migration *in vitro*. Fibroblasts play a key role in tissue repair by producing extracellular matrix components and promoting wound closure. Stimulating their migration is crucial in regenerative medicine since it could help to treat wound healing, support angiogenesis, and reduces fibrosis, ultimately improving tissue regeneration.

In conclusion, the derivation of iMSCs from AFSC-iPSCs addresses critical limitations of primary AFSCs, including donor variability and replicative senescence, which constrain their scalability for clinical applications. These findings underscore the potential of iMSCs as a robust and reproducible source of therapeutic cells, likely attributed to the epigenetic reprogramming of the AFSC genome, offering a promising strategy to overcome the challenges associated with primary AFSCs.

## References

1. Abdulrazzak, H., Moschidou, D., Jones, G., and Guillot, P.V. (2010). Biological characteristics of stem cells from foetal, cord blood and extraembryonic tissues. J R Soc Interface 7 *Suppl 6*, S689–706. 10.1098/rsif.2010.0347.focus.

2. Jones, G.N., Moschidou, D., Puga-Iglesias, T.-I., Kuleszewicz, K., Vanleene, M., Shefelbine, S.J., Bou-Gharios, G., Fisk, N.M., David, A.L., Coppi, P.D., et al. (2012). Ontological differences in first compared to third trimester human fetal placental chorionic stem cells. PLoS One 7, e43395.

3. Coppi, P.D., Bartsch, G.J., Siddiqui, M.M., Xu, T., Santos, C.C., Perin, L., Mostoslavsky, G., Serre, A.C., Snyder, E.Y., Yoo, J.J., et al. (2007). Isolation of amniotic stem cell lines with potential for therapy. Nat Biotechnol 25, 100–106. 10.1038/nbt1274.

4. Loukogeorgakis, S.P., and Coppi, P.D. (2017). Concise Review: Amniotic Fluid Stem Cells: The Known, the Unknown, and Potential Regenerative Medicine Applications. Stem Cells 35, 1663–1673. 10.1002/stem.2553.

5. Guillot, P.V., O’Donoghue, K., Kurata, H., and Fisk, N.M. (2006). Fetal stem cells: betwixt and between. Semin Reprod Med 24, 340–347. 10.1055/s-2006-952149.

6. Guillot, P.V., Bari, C.D., Dell’Accio, F., Kurata, H., Polak, J., and Fisk, N.M. (2008). Comparative osteogenic transcription profiling of various fetal and adult mesenchymal stem cell sources. Differentiation 76, 946–957. 10.1111/j.1432-0436.2008.00279.x.

7. Guillot, P.V., Gotherstrom, C., Chan, J., Kurata, H., and Fisk, N.M. (2007). Human first-trimester fetal MSC express pluripotency markers and grow faster and have longer telomeres than adult MSC. Stem Cells 25, 646–654. 10.1634/stemcells.2006-0208.

8. Moschidou, D., Corcelli, M., Hau, K.-L., Ekwalla, V.J., Behmoaras, J.V., Coppi, P.D., David, A.L., Bou-Gharios, G., Cook, H.T., Pusey, C.D., et al. (2016). Human Chorionic Stem Cells: Podocyte Differentiation and Potential for the Treatment of Alport Syndrome. Stem Cells Dev 25, 395–404. 10.1089/scd.2015.0305.

9. Vanleene, M., Porter, A., Guillot, P.-V., Boyde, A., Oyen, M., and Shefelbine, S. (2012). Ultra-structural defects cause low bone matrix stiffness despite high mineralization in osteogenesis imperfecta mice. Bone 50, 1317–1323. 10.1016/j.bone.2012.03.007.

10. Vanleene, M., Saldanha, Z., Cloyd, K.L., Jell, G., Bou-Gharios, G., Bassett, J.H.D., Williams, G.R., Fisk, N.M., Oyen, M.L., Stevens, M.M., et al. (2011). Transplantation of human fetal blood stem cells in the osteogenesis imperfecta mouse leads to improvement in multiscale tissue properties. Blood 117, 1053–1060. 10.1182/blood-2010-05-287565.

11. Jones, G.N., Moschidou, D., Abdulrazzak, H., Kalirai, B.S., Vanleene, M., Osatis, S., Shefelbine, S.J., Horwood, N.J., Marenzana, M., Coppi, P.D., et al. (2014). Potential of human fetal chorionic stem cells for the treatment of osteogenesis imperfecta. Stem Cells Dev 23, 262–276. 10.1089/scd.2013.0132.

12. Guillot, P.V., Abass, O., Bassett, J.H.D., Shefelbine, S.J., Bou-Gharios, G., Chan, J., Kurata, H., Williams, G.R., Polak, J., and Fisk, N.M. (2008). Intrauterine transplantation of human fetal mesenchymal stem cells from first-trimester blood repairs bone and reduces fractures in osteogenesis imperfecta mice. Blood 111, 1717–1725. 10.1182/blood-2007-08-105809.

13. Ranzoni, A.M., Corcelli, M., Hau, K.-L., Kerns, J.G., Vanleene, M., Shefelbine, S., Jones, G.N., Moschidou, D., Dala-Ali, B., Goodship, A.E., et al. (2016). Counteracting bone fragility with human amniotic mesenchymal stem cells. Sci Rep 6, 39656. 10.1038/srep39656.

14. Corcelli, M., Hawkins, K., Vlahova, F., Hunjan, A., Dowding, K., Coppi, P.D., David, A.L., Peebles, D., Gressens, P., Hagberg, H., et al. (2018). Neuroprotection of the hypoxic-ischemic mouse brain by human CD117(+)CD90(+)CD105(+) amniotic fluid stem cells. Sci Rep 8, 2425. 10.1038/s41598-018-20710-9.

15. Sagar, R.L., Åström, E., Chitty, L.S., Crowe, B., David, A.L., DeVile, C., Forsmark, A., Franzen, V., Hermeren, G., Hill, M., et al. (2024). An exploratory open-label multicentre phase I/II trial evaluating the safety and efficacy of postnatal or prenatal and postnatal administration of allogeneic expanded fetal mesenchymal stem cells for the treatment of severe osteogenesis imperfecta in infants and fetuses: the BOOSTB4 trial protocol. BMJ Open 14, e079767. 10.1136/bmjopen-2023-079767.

16. Chan, J., Kumar, S., Fisk, N.M., Sagar, R., Walther-Jallow, L., David, A.L., Hill, M., Lewis, C., Riddington, M., Crowe, B., et al. (2019). Stakeholder views and attitudes towards prenatal and postnatal transplantation of fetal mesenchymal stem cells to treat Osteogenesis Imperfecta. Eur J Hum Genet 27, 1244–1253. 10.1038/s41431-019-0387-4.

17. Moschidou, D., Drews, K., Eddaoudi, A., Adjaye, J., Coppi, P.D., and Guillot, P.V. (2013). Molecular signature of human amniotic fluid stem cells during fetal development. Curr Stem Cell Res Ther 8, 73–81.

18. Hawkins, K.E., Moschidou, D., Faccenda, D., Wruck, W., Martin-Trujillo, A., Hau, K.-L., Ranzoni, A.M., Sanchez-Freire, V., Tommasini, F., Eaton, S., et al. (2017). Human Amniocytes Are Receptive to Chemically Induced Reprogramming to Pluripotency. Mol Ther 25, 427–442. 10.1016/j.ymthe.2016.11.014.

19. Moschidou, D., Mukherjee, S., Blundell, M.P., Drews, K., Jones, G.N., Abdulrazzak, H., Nowakowska, B., Phoolchund, A., Lay, K., Ramasamy, T.S., et al. (2012). Valproic Acid Confers Functional Pluripotency to Human Amniotic Fluid Stem Cells in a Transgene-free Approach. Mol. Ther. 20, 1953–1967. 10.1038/mt.2012.117.

20. Moschidou, D., Mukherjee, S., Blundell, M.P., Jones, G.N., Atala, A.J., Thrasher, A.J., Fisk, N.M., Coppi, P.D., and Guillot, P.V. (2013). Human mid-trimester amniotic fluid stem cells cultured under embryonic stem cell conditions with valproic acid acquire pluripotent characteristics. Stem Cells Dev 22, 444–458. 10.1089/scd.2012.0267.

21. Peister, A., Deutsch, E.R., Kolambkar, Y., Hutmacher, D.W., and Guldberg, R.E. (2009). Amniotic Fluid Stem Cells Produce Robust Mineral Deposits on Biodegradable Scaffolds. Tissue Eng. Part A 15, 3129–3138. 10.1089/ten.tea.2008.0536.

22. Sun, H., Feng, K., Hu, J., Soker, S., Atala, A., and Ma, P.X. (2010). Osteogenic differentiation of human amniotic fluid-derived stem cells induced by bone morphogenetic protein-7 and enhanced by nanofibrous scaffolds. Biomaterials 31, 1133–1139. 10.1016/j.biomaterials.2009.10.030.

23. Trapani, M.D., Bassi, G., Fontana, E., Giacomello, L., Pozzobon, M., Guillot, P.V., Coppi, P.D., and Krampera, M. (2015). Immune regulatory properties of CD117(pos) amniotic fluid stem cells vary according to gestational age. Stem Cells Dev 24, 132–143. 10.1089/scd.2014.0234.

24. Zare, E., Hosseini, E.S., Azad, F.S., Javid, A., Javazm, R.R., Abessi, P., Montazeri, F., and Hoseini, S.M. (2025). Replicative senescence in amniotic fluid-derived mesenchymal stem cells and its impact on their immunomodulatory properties. Histochem. Cell Biol. 163, 34. 10.1007/s00418-025-02364-7.

25. Chen, Y.S., Pelekanos, R.A., Ellis, R.L., Horne, R., Wolvetang, E.J., and Fisk, N.M. (2012). Small molecule mesengenic induction of human induced pluripotent stem cells to generate mesenchymal stem/stromal cells. Stem Cells Transl Med 1, 83–95. 10.5966/sctm.2011-0022.

26. Wruck, W., Graffmann, N., Spitzhorn, L.-S., and Adjaye, J. (2021). Human Induced Pluripotent Stem Cell-Derived Mesenchymal Stem Cells Acquire Rejuvenation and Reduced Heterogeneity. Front. Cell Dev. Biol. 9, 717772. 10.3389/fcell.2021.717772.

27. Jungbluth, P., Spitzhorn, L.-S., Grassmann, J., Tanner, S., Latz, D., Rahman, M.S., Bohndorf, M., Wruck, W., Sager, M., Grotheer, V., et al. (2019). Human iPSC-derived iMSCs improve bone regeneration in mini-pigs. Bone Res. 7, 32. 10.1038/s41413-019-0069-4.

28. Spitzhorn, L.-S., Megges, M., Wruck, W., Rahman, M.S., Otte, J., Degistirici, Ö., Meisel, R., Sorg, R.V., Oreffo, R.O.C., and Adjaye, J. (2019). Human iPSC-derived MSCs (iMSCs) from aged individuals acquire a rejuvenation signature. Stem Cell Res. Ther. 10, 100. 10.1186/s13287-019-1209-x.

29. Hass, R., Kasper, C., Böhm, S., and Jacobs, R. (2011). Different populations and sources of human mesenchymal stem cells (MSC): A comparison of adult and neonatal tissue-derived MSC. Cell Commun. Signal. 9, 12. 10.1186/1478-811x-9-12.

